# Mapping the probability of forest snow disturbances in Finland

**DOI:** 10.1101/2020.12.23.424139

**Authors:** S Suvanto, A Lehtonen, S Nevalainen, I Lehtonen, H Viiri, M Strandström, M Peltoniemi

## Abstract

The changing forest disturbance regimes emphasize the need for improved damage risk information. Here, our aim was to (1) improve the current understanding of snow damage risks by assessing the importance of abiotic factors, particularly the modelled snow load on trees, versus forest properties in predicting the probability of snow damage, (2) produce a snow damage probability map for Finland. We also compared the results for winters with typical snow load conditions and a winter with exceptionally heavy snow loads. To do this, we used damage observations from the Finnish national forest inventory (NFI) to create a statistical snow damage occurrence model, spatial data layers from different sources to use the model to predict the damage probability for the whole country in 16 x 16 m resolution. Snow damage reports from forest owners were used for testing the final map. Our results showed that best results were obtained when both abiotic and forest variables were included in the model. However, in the case of the high snow load winter, the model with only abiotic predictors performed nearly as well as the full model and the ability of the models to identify the snow damaged stands was higher than in other years. The results showed patterns of forest adaptation to high snow loads, as spruce stands in the north were less susceptible to damage than in southern areas and long-term snow load reduced the damage probability. The model and the derived wall-to-wall map were able to discriminate damage from no-damage cases on a good level. The damage probability mapping approach identifies the drivers of snow disturbances across forest landscapes and can be used to spatially estimate the current and future disturbance risks in forests, informing practical forestry and decision-making and supporting the adaptation to the changing disturbance regimes.

## Introduction

Forest disturbances caused by snow are frequent in high latitude regions (Valinger and Fridman, 1997; Jalkanen and Mattila, 2000; Díaz-Yáñez et al., 2016, Duperat et al. 2020) and high-altitude areas (Klopcic et al., 2009; Hlásny et al., 2011; Nagel et al., 2017). In Europe, the estimates of forest damage caused by snow disturbance events range from 1 to 4 million m3 of wood per year (Nykänen et al., 1997; Schelhaas et al., 2003). While climate warming may lead to reduced levels of snow disturbances (Seidl et al., 2017) the future changes are likely to be spatially asymmetric. For example, snow damage is projected to decrease in southern and western Finland but in northern and eastern parts of the country heavy snow loads are expected to increase. This is because the warmer and more humid climate will increase the occurrence of wet snow hazard events and conditions favorable for rime accumulation in these areas (Lehtonen et al., 2016; Venäläinen et al., 2020).

Snow disturbances are an inherent part of the forest ecosystem in northern and high-altitude forests. They cause economic losses in terms of damaged wood and increased tree mortality. Snow disturbances in forests also damage the infrastructure; the power grid in particular is vulnerable as tree tops and trees with heavy crown snow loads fall on the power lines. Snow damaged trees and areas are also more susceptible to subsequent damage by insects or fungi (Nykänen et al., 1997). Many of the negative effects of snow disturbances could potentially be alleviated by improved planning and forest management, but this requires accurate information about the damage risks. Spatial risk information is increasingly required by the society and it is used actively in management, operations and financial planning among owners, industry, and insurers.

Precise forest and climate data has made it possible to present risk information at high resolution. For example, Suvanto et al. (2019), mapped forest wind damage probabilities in forests at 16 m x 16 m resolution, by using a model that drew from damage observations made in the Finnish national forest inventory (NFI), spatially identified high wind areas, and environmental and forest resource data from various open data sources. The high spatial resolution of the map allows the consideration of disturbance probability on the level of individual forest stands, i.e. the spatial unit in which the management decisions are being made. In northern forests, snow disturbances play an important role, and therefore a better understanding of how snow damage risks can be predicted at large scale but at high resolution is needed.

Snow damage to trees is induced when the forces generated by a large crown snow load, often together with wind, exceed the force required to break the stem of the tree. Meteorological data is crucial in modelling forest snow disturbances, as specific meteorological conditions are needed for snow to accumulate on trees. Typically snow accumulates to trees typically within a narrow temperature range close to 0 °C (Solantie, 1994). Conditions after the snowfall are important for the damage, as retention of snow in the tree crowns is temperature dependent (Nykänen et al., 1997). As the accumulation of rime and snow on trees is driven by temperature and wind conditions, topographic factors are typically correlated to the occurrence of snow damage (Nykänen et al., 1997, Lehtonen et al., 2014). Snow load on trees can be categorized in different types, such as rime, wet snow, dry snow and frozen snow, and the physical process of snow accumulation differs by the type. Lehtonen et al. (2014) showed that improved results in modelling snow load in tree crowns could be achieved by considering the different snow load types separately.

The characteristics of the forest stand and the trees play an important role, as damage occurs when the gravitational forces and torque caused by the crown snow load exceed the stem tolerance limit. The tolerance is largely related to stem taper and characteristics of the tree crown, while these are driven by factors such as tree species and stand characteristics (Nykänen et al., 1997; Peltola et al., 1999). From a biomechanical perspective, older trees with stronger stem taper and thicker stems should be more resistant to crown snow loads than smaller trees with modest stem taper and thinner stems. The density of stand may indirectly affect the susceptibility of trees to damage, as density-driven competition drives the growth of thin and tall stems (Nykänen et al., 1997; Peltola et al., 1999).

Coniferous species are generally more susceptible to damage than deciduous trees, and Norway spruce is less vulnerable compared to Scots pine (Jalkanen and Konocpka, 1998; Nykänen et al., 1997). Tree structural properties predisposing trees to damage vary also within species. In Norway spruce, the tree morphology varies across the species range so that in high altitude and latitude areas the narrow crown shape and dense, horizontal branches reduce the accumulation of snow on the crowns, decreasing the probability of snow damage (Mikola, 1938, Morgenstern 1996, Geburek et al., 2008).

In this study, our aim was to (1) assess the importance of meteorological and topographic factors versus forest properties for the occurrence probability of snow damage in forests, comparing results from winters with typical snow load conditions and an exceptionally heavy snow load winter, and (2) produce a snow damage probability map for Finland and test the ability of the map to identify the stands vulnerable to snow disturbances. As the meteorological variable, we used model-derived crown snow load, which should be the best proxy for damage-causing climatic conditions and which allows predicting snow damage risks changes under climate change conditions.

## Material and methods

### National forest inventory data

National forest inventory (NFI) data was used for the snow damage observations and for the forest characteristics data. The used data included plots from the 10th (2005-2008), 11th (2009-2013) and 12th (2014-2018) Finnish NFIs (Korhonen et al. 2016, 2017). NFI10 measurements from 2004 were excluded as no full 5 year period of snow load data was available before that year. To avoid having repeated measurements from the same plots in the data, only temporary NFI plots from NFI10 and NFI11 were included in the analysis, whereas all plots (temporary and permanent) were included from the NFI12. Only NFI plots on forest land were included and plots on treeless stands were excluded from the data. Data points with missing data in any of the used predictor variables were excluded in the analysis. The final data consisted of a total 111 677 plots, in 2 380 of which snow damage was recorded (Table 1).

**Table 1.** Statistics of stand level snow damage, damage severity and damage type in the NFI data.

|  | All | 2005-2017 | 2018 |
| --- | --- | --- | --- |
| Total number of plots | 111 677 | 102 671 | 9 006 |
| Total damaged plots | 2 380 | 1 885 | 495 |
| % damaged plots | 2.13 | 1.84 | 5.50 |
| <u>Damage severity (% of cases)</u> |  |  |  |
| 0, slight damage | 57.4 | 59.4 | 49.9 |
| 1, moderate damage | 38.9 | 37.3 | 44.8 |
| 2, severe damage | 3.7 | 3.2 | 5.3 |
| <u>Damage type (% of cases)</u> |  |  |  |
| Dead standing trees | 0.5 | 0.5 | 0.6 |
| Uprooted or broken trees | 75.8 | 75.6 | 76.4 |
| Stem damage | 0.5 | 0.5 | 0.4 |
| Dead or broken crowns | 12.3 | 10.6 | 18.8 |
| Other crown damage | 10.7 | 12.5 | 3.6 |
| Branch damage | 0.2 | 0.2 | 0.2 |
| Defoliation | <0.1 | 0.1 | -- |
| Discolouration | <0.1 | 0.1 | -- |

Stand level snow damage observations from the Finnish national forest inventory (NFI) were used in the study. All damage cases that occurred in the dominant tree storey of the stand (i.e., the tree storey that determines silvicultural operations for the stand) and where the causal agent of the primary damage had been classified as “snow” and the timing of the damage was estimated to be within 5 years were included as damage in the analysis.

The damage type was most often fallen or broken trees (no distinction of these two are made in the data) but also other damage types were found (Table 1). Damage severity is recorded in the NFI as cumulative effect of all damage agents found in the stand, and no information about the severity of snow damage specifically is included if also other damage causes were present. Severity is assessed on a four-point scale (0 to 3) and most stands with snow damage are classified to the two lowest classes (0 = modest damage, does not affect the silvicultural quality of the stand or change the development class, and 1 = moderate damage, lowers the silvicultural quality of the stand by one class), with some observations in class second highest class (2 = severe damage, decreases the quality of the stand by more than one class) and no observations in the highest damage severity class (3 = complete damage, immediate regeneration required; Table 1).

Other information from the NFI used in our analysis included stand dominant tree species, average tree height and diameter at breast height (DBH) in stand, basal area, forest management operations (thinning, tending of seedling stands) and their timing, site type and proportions of basal area represented by different species. From the species data we derived variables describing the total number of tree species in the plot, proportion of basal area covered by the species with the highest basal area and the Shannon diversity index, which was also calculated from shares of basal area for each species.

Stand average DBH was not recorded for stands of development class “young seedling stand”, where the height of the dominant tree species is less than 1.3 meters. For these stands DBH was set to 0 cm. In NFI10, DBH was also missing for the development class “advanced seedling stand”. For these, the DBH was estimated based on the measurements in NFI11 and NF12. DBH in this development class was predicted based on average tree height and dominant tree species by fitting a GLM model with gamma distribution and log-link function to the NFI11 and NFI12 data where the DBH was available, and then using this model to predict the DBH values for the advanced seedling stands in NFI10 where the DBH information was missing.

### Snow load on trees

Maximum snow load on tree canopies was calculated for each winter for years 2001 to 2018, using the snow load model of the Finnish Meteorological Institute (FMI) (Lehtonen et al., 2014) and the ERA5 reanalysis data (Hoffmann et al., 2019).

The snow-load model is a statistical model in operational use at the FMI. The model assumes a tree with cone-shaped crown with a projected catchment area of one square meter from above and from the side in the direction of the wind and calculates the snow load on tree canopies in four different snow accumulation types: rime, dry snow, wet snow and frozen snow (Lehtonen et al., 2014). Here, the sum of the different snow load types was used, and the maximum snow load of the previous five years from the NFI measurement date was used for each NFI plot, as the snow damage observed on the plots may have occurred within 5 previous years.

### Topographic variables

Altitude as meters above sea level was extracted for the NFI plot locations from the 25 meter resolution digital elevation model (DEM) from the National Survey of Finland. Relative elevation was calculated from the same DEM as the difference between the altitude at the plot location and the average altitude within 1 kilometer radius. Thus, negative values of the variable represent with topographic positions lower than the near surroundings and positive values higher.

### Statistical modelling

Statistical models were fitted using the occurrence of snow damage in the NFI plots as the binary response variable and forest properties, snow load data and topographic variables as predictors. Only snow damage cases that had occurred within 5 years of the NFI field measurement date (according to the estimate of the field team) were considered.

Two different types of statistical modelling methods were used: generalized linear models (GLM) and generalized additive models (GAM), both with a logistic link function. GAM is an extension of a GLM where the linear predictor contains a sum of smooth functions of continuous predictors. Using smooth functions instead of detailed parametric relationships (as done in GLMS) allows for more flexibility in the dependence of the response of the predictors (Wood, 2017).

The model selection was done using only the GLM model. The model predictors were chosen based on (1) existing understanding of snow damage dynamics, (2) availability of national extent GIS-data to be used for map prediction, (3) statistical significance of highest order terms in model, requiring significance on the level of p < 0.001, as the large sample size easily leads to small p-values, (4) improvement in AIC when comparing alternative models and (5) collinearity between predictors, determined by the generalized variation inflation factor (GVIF). If the GVIF exceeded 4 for any of the predictor variables, one of the correlated variables was left out of the model. The decision on which variable to exclude was made following the same five steps of comparing alternative models. For continuous variables with non-negative values, log-transformations with natural logarithm were tested and included where they led to a lower AIC. For transparency of the model selection process, intermediate model versions with variables not included in the final model can be found in the supplementary material (S1).

The potential predictor variables considered in the model selection were grouped into abiotic variables relating to snow load and topography (ABIOTIC) and forest variables (FOREST). The ABIOTIC variable group contained variables describing crown snow load (maximum of previous 5 years), long term average of winter maximum crown snow load, altitude from sea level, relative elevation in comparison to a kilometer radius and a variable describing if the plot was located in the north boreal vegetation zone, according to the biogeographical zones data from the Finnish Environment Institute (SYKE 2015). The FOREST variable group included dominant tree species of the stand, average DBH of the stand, average tree height of the stand, basal area, forest management history, site type (poor vs fertile, using the same classification as in Suvanto et al. 2019), number of tree species, proportion of basal area by the most abundant species and the Shannon diversity index, calculated from the proportions of basal area by each species. For forest management history, three variables were included – all thinnings, pre-commercial thinnings and tending of seedling stands. All were included as presence/absence variables that described if the management operation had been carried out at the stand more than 5 years ago. Management information within five years from the NFI measurement was not considered because, if snow damage had occurred in the stand, it would not be clear if the management was done before or after damage (damage was considered from the latest 5 years). To find potential species specific responses, interaction terms were tested between tree species and DBH, basal area, the snow load variables and the north boreal zone variable.

After the predictors were selected for the model, two additional submodels were formed to have three models: a full model with all predictors (FULL), a model with only abiotic predictors (ABIOTIC) and a model with only predictors related to forest properties (FOREST) (see variables included in each group in the final model in Table 3, results for the variables not included in the final model can be found in S1). In case of an interaction between variables in different variables groups, both variables were included in the FOREST group.

**Table 2.**
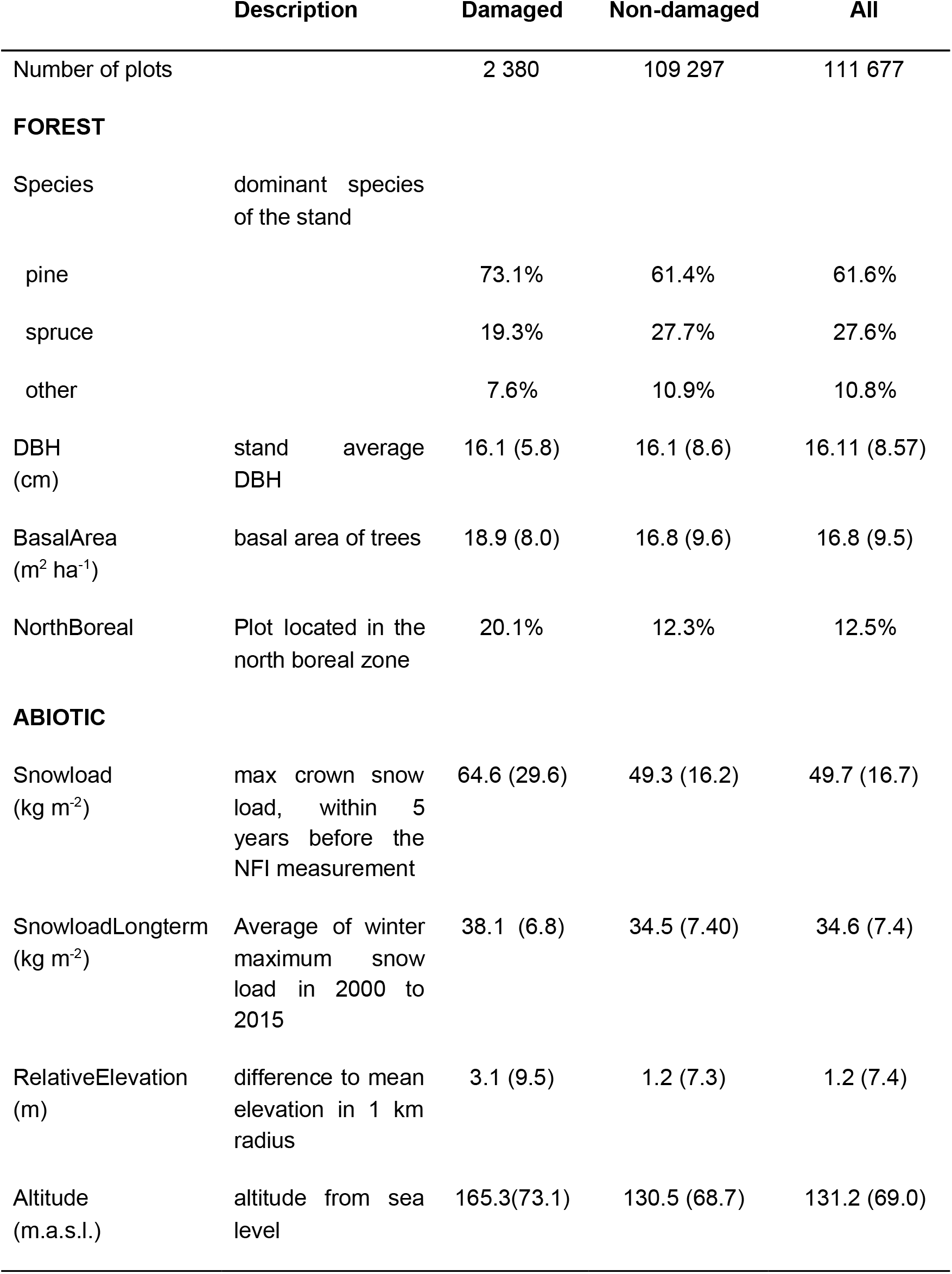
Number of plots and the descriptive statistics for forest, topographical and snow load variables for damaged and non-damaged plots separately and for all the plots in the data. For categorical variables values represent percentages of plots in each class and for continuous variables mean and standard deviation, the latter in parenthesis.

**Table 3.** Model results for the full GLM model

|  | **Estimate** | **Std. Error** | **z value** | **Pr(>\|z\|)** |
| --- | --- | --- | --- | --- |
| Intercept | -7.209 | 0.168 | -42.975 | < 0.001 |
| SpeciesSpruce<sup>1)</sup> | -0.287 | 0.058 | -4.952 | < 0.001 |
| SpeciesOther<sup>1)</sup> | 0.716 | 0.214 | 3.350 | < 0.001 |
| DBH | -0.072 | 0.004 | -16.308 | < 0.001 |
| log(Basalarea + 0.5) | 1.101 | 0.048 | 22.721 | < 0.001 |
| NorthBoreal | -0.031 | 0.069 | -0.450 | < 0.001 |
| SnowloadLongterm | -0.027 | 0.005 | -6.093 | < 0.001 |
| Snowload | 0.032 | 0.001 | 30.822 | < 0.001 |
| RelativeElevation | 0.026 | 0.002 | 10.752 | < 0.001 |
| Altitude | 0.006 | 4.7E-04 | 11.892 | < 0.001 |
| SpeciesOther x DBH | -0.092 | 0.016 | -5.670 | < 0.001 |
| SpeciesSpruce x NorthBoreal | -0.749 | 0.186 | -4.029 | < 0.001 |

<sup>1)</sup> Compared to the reference species Scots pine

Models with the same predictor variables were then fitted as generalized additive models (GAM) to test if using a non-parametric model would lead to better outcome, as they are able to effectively deal with non-linear relationships. Continuous predictor variables were included in the GAM models as smoothing spline functions. The dimension parameter (k), that sets the upper limit on the degrees of freedom related to the smooth, was set to 15 for all variables. The suitability of the k parameter was assessed visually. In addition, the effective degrees of freedom after fitting the model were lower than k for all of the terms, suggesting that the chosen k values were sufficiently large.

The performance of the models was assessed with 10-fold stratified cross-validation, where the number of damaged plots was divided evenly into the folds. One fold at the time was used as test data while the model was fitted with the remaining nine folds. Receiver operating characteristic (ROC) and area under curve (AUC) were calculated for the test data to assess the model performance. AUC value of 0.5 corresponds to a situation where the model does not do better than randomly assigning the prediction values whereas AUC value of 1 would mean that the model is perfectly able to discriminate between damage cases and no-damage cases. As a rule of thumb, 0.7 is often used as an acceptable level of discrimination between the classes (Hosmer et al., 2013).

To compare the results for typical snow load winters and an exceptionally high snow load winter, AUC values for the cross-validation were calculated in three different subsets: using all the data in the test data fold, using only data from 2005-2017 in the test data fold (“typical snow load winters”) and using only data from the 2017-2018 winter (“exceptional snow load winter”, Fig. 1). These subsets were only used in the test data fold, all data in the remaining folds were used to fit the model in each cross-validation round.

**Figure 1.**
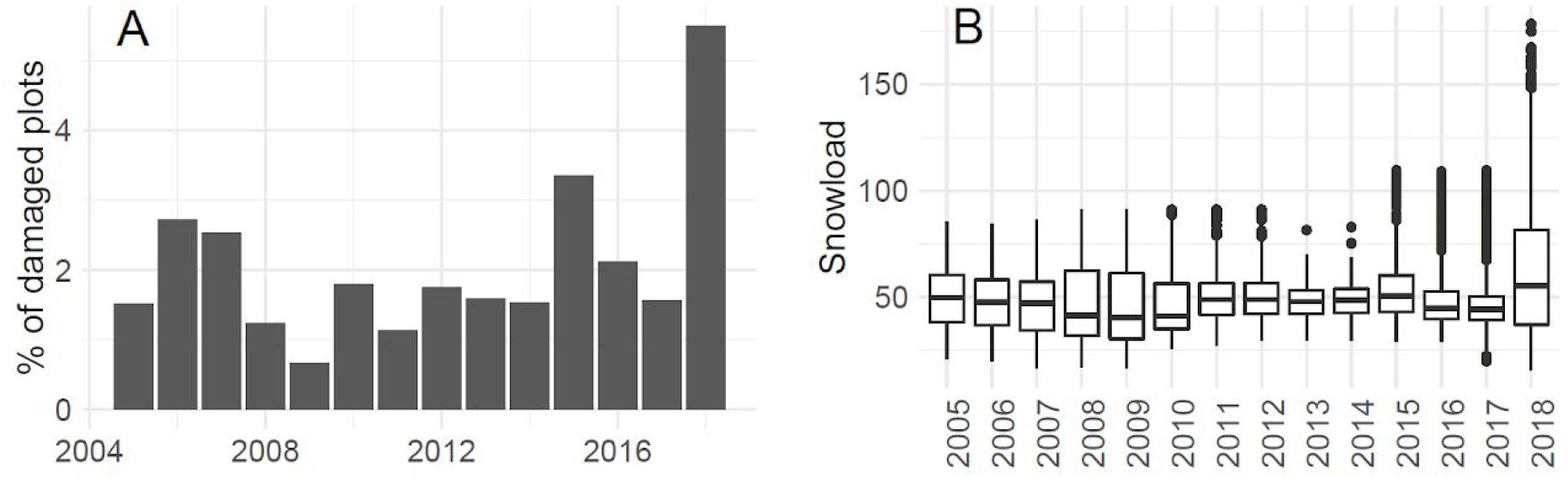
Percentage of plots with snow damage in each year (A; year refers to the year the NFI plot has been measured on the field, damage may have occurred within previous five years) and (B) maximum snow load at the NFI plots within a five year time window.

Statistical modelling was done in R version 3.5.2., ROC and AUC were calculated with the R package *pROC* (Robin et al., 2011). The GAMs were fitted using the R package *mgcv* (Wood, 2017).

### Mapping of damage probability

The snow damage probability map was calculated for the whole country of Finland in 16 x 16 m pixel resolution, by using the full GLM and GAM models and geographic information system (GIS) datasets representing the predictors of models.

Regarding GIS datasets, multi-source forest inventory (MS-NFI) forest resource maps for 2017 (Mäkisara et al., 2019) were used for the forest variables (tree species, DBH, basal area). Topographic variables (altitude and relative elevation) were derived from the 25 meter resolution DEM of the National Land Survey of Finland and resampled to the same 16 m x 16 m grid. Snow load data (Lehtonen et al. 2014) for winter 2017-2018 was used in the calculation of the map, as this winter was also used for the testing of the map.

The processing of GIS data was conducted using R (package raster), Python and GDAL. The calculation of the map was done using R package raster (Hijmans, 2017) and the sp package (Pebesma and Bivand, 2005).

### Testing the map

The test data for the damage risk maps for winter 2017-2018 was obtained from the Finnish Forest Centre.

For damage events, forest use declarations where snow damage had been recorded were extracted from the data, using the reports sent to the Forest Centre from December 1st 2017 to September 30th 2018. Forest owners are required by law to submit a forest use declaration to the Forest Centre before conducting forest management operations at their stands and since 2012 these declarations have included information about forest damage in the stand in case the damage has been the reason for the logging operation. The declarations contain information about the stand, including the occurred damage, and a spatially referenced polygon outlining the stand. The final test data contained a total of 11 807 snow damaged stands (referred to as “snow damage polygons” from now on).

To compare the snow damage polygons from forest use declarations to non-damaged stands, we used another data set by the Forest Centre, that contains spatial polygons and basic forest property information for forests on private lands in Finland. From this data, one percent of the polygons in the whole country was randomly sampled. Polygons classified as open stands (i.e., did not have trees) were excluded from the sample. While this data set does not contain information about forest damage, we assume that these stands are not damaged. The resulting data consisted of 101 073 polygons (referred as “non-damaged polygons” from now on).

To test if the map was able to differentiate between damaged and non-damaged stands within the larger damage area (as compared to only differentiating the general damage area from the rest of the country), another test was carried out by only including the non-damaged polygons that were located within 10 kilometers from the damaged stands (Fig. 5). This subset contained 16 486 non-damaged polygons.

For both snow damage polygons and non-damaged polygons the average value of snow damage map pixels within each polygon was calculated for both maps based on GLM and GAM models. Then, the distribution of the map values was examined on the snow damaged and non-damaged maps, and ROC curves and AUC values were calculated to assess the performance of the maps to identify the snow damage cases.

Both of the used data sets (forest use declarations and stand polygons for private lands) are published by the Finnish Forest Centre under CC BY 4.0 licence and are openly available (https://www.metsaan.fi/paikkatietoaineistot). Data were loaded in October 2020.

## Results

Both GLM model results show that abiotic factors, especially crown snow load, drive the snow damage, as damage probability increases with increasing snow load, relative elevation and altitude (Table 3, Fig. 2). Yet, forest characteristics also have an impact on damage occurrence. Damage probability was higher in stands with higher basal area and in stands with lower average DBH. The model showed higher damage probabilities in stands dominated by pine compared to other species. Norway spruce dominated stands show regional different patterns, with disturbance probability being significantly lower in the north boreal zone compared to other parts of the country. For species group “other”, mainly consisting of birches, higher values of damage probability were predicted for small DBH stands compared to pine and spruce (Fig. 2).

**Figure 2.**
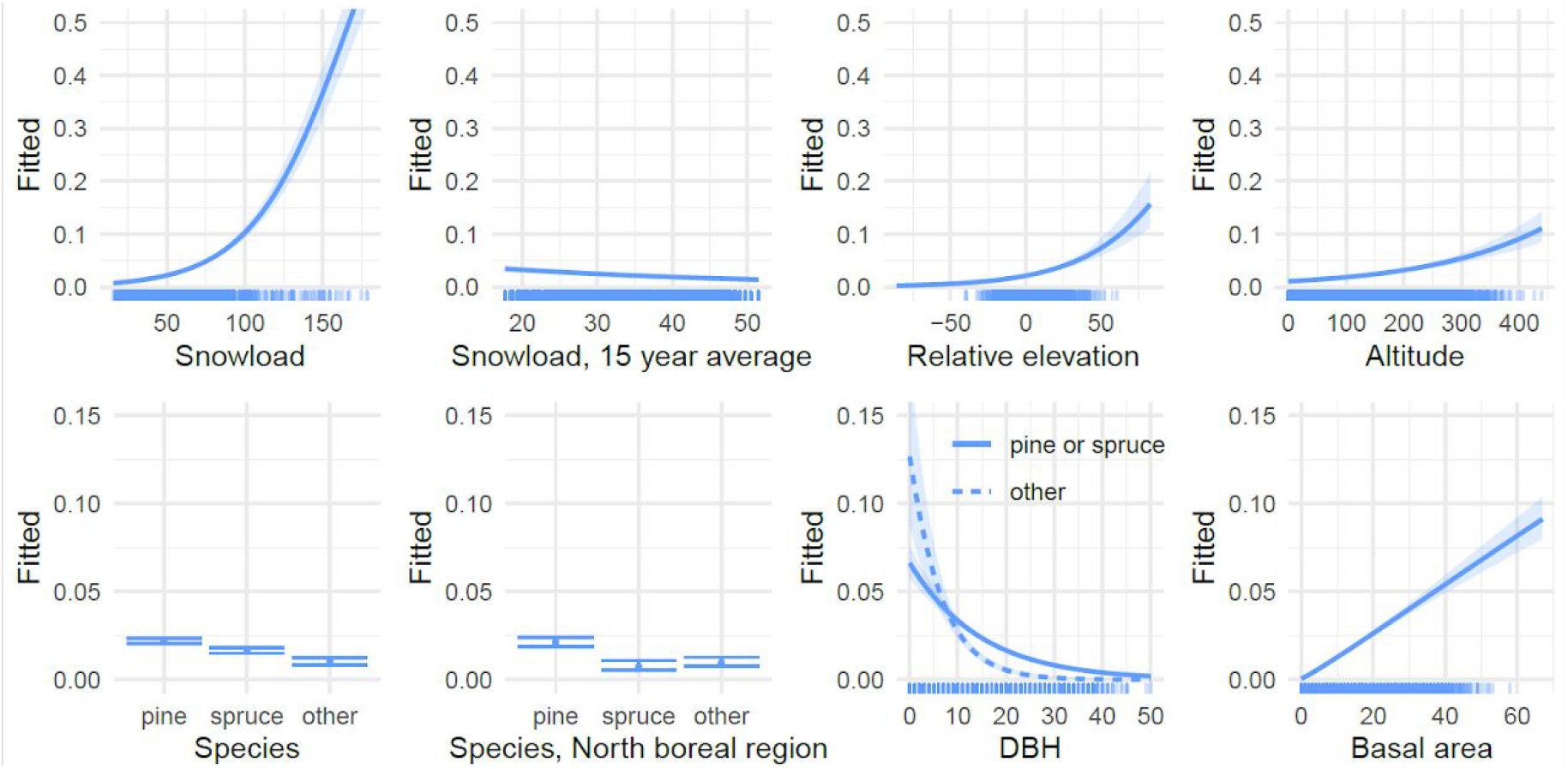
The impact of predictors for the probability of snow damage occurrence according to the full GLM model. Note different y-axis limits in abiotic variables (upper row) and the forest variables (lower row). The rug showing the distribution of data points is a random subset of 10 000 plots from the original data.

GAM models showed generally similar patterns as GLM models but also revealed non-linearities not visible in the GLM results. For example, probability of damage only started to rise drastically with snow load after 75 kg m^-2^ (Fig. 3), which is clearly higher than the snow loads observed in typical winter conditions (Fig. 1B). The GAM results also show decrease of damage probability with relative elevation and altitude after a certain thresholds, but as there are few observations at high values of both of these variables, there is high uncertainty of the shape of the spline. Long term snow load (15 years average) also showed a nonlinear trend with the damage probability, with damage probability values peaking at 30 kg m^2^.

**Figure 3.**
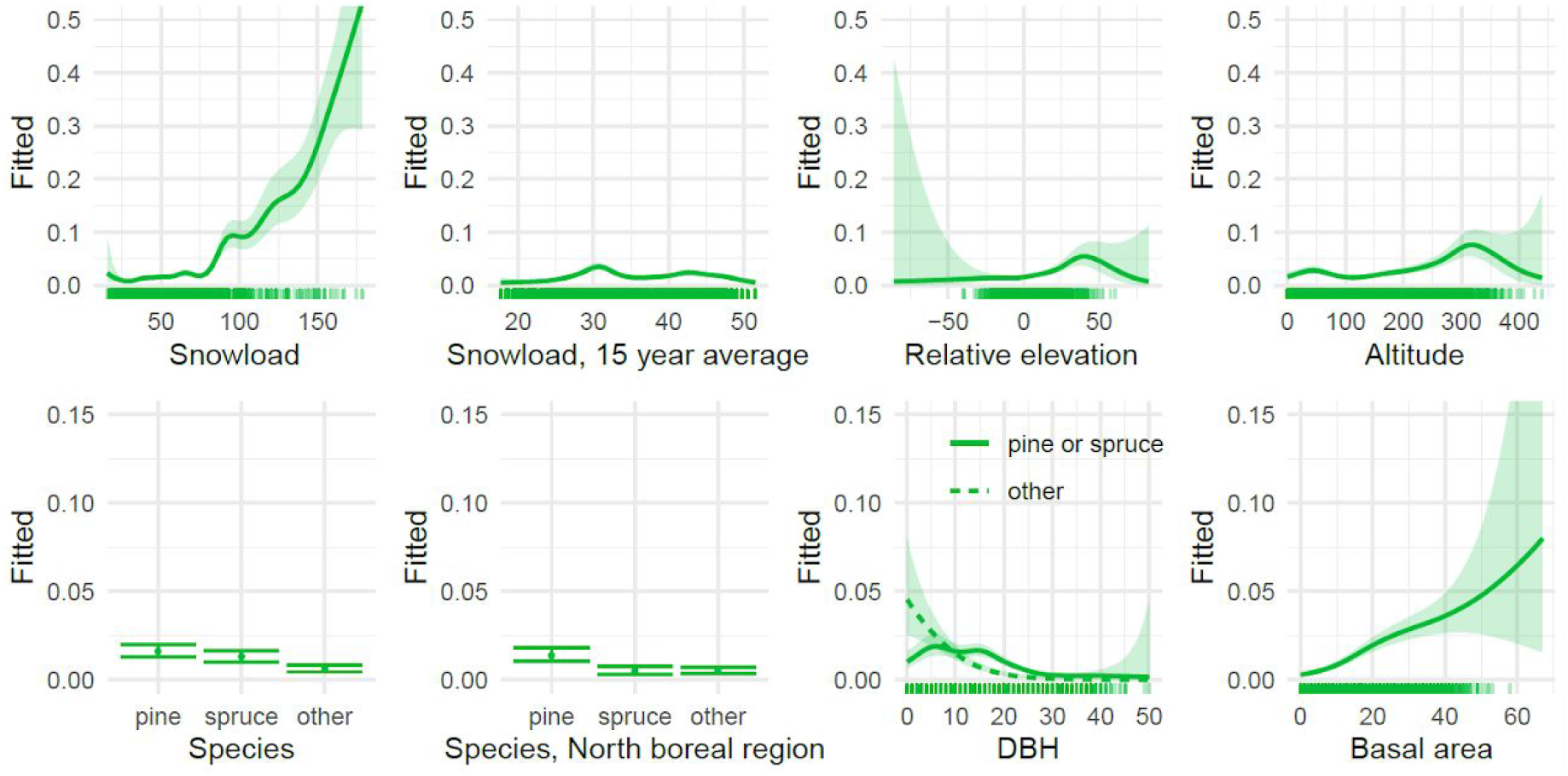
Effects plots for predictors in the full GAM model. Note different y-axis limits in abiotic variables (upper row) and the forest variables (lower row). The rug showing the distribution of data points is a random subset of 10 000 plots from the original data.

The forest management variables (thinnings, precommercial thinning and tending of seedling stands) were not included in the final model as the p-values of the coefficients were larger than the defined p < 0.001 level. For tending of seedling stands the p-values were rather close to this level (p = 0.0018), but the variable was nevertheless excluded for not meeting the set criteria and also for the difficulty of attaining relevant GIS data (results in supplementary material S1). Similarly, the species composition variables were excluded from the model, with results for Shannon diversity index being closest for being included (p = 0.0041) showing negative effect on damage probability (S1).

The cross-validation of the models showed that the FULL model with both abiotic and forest variables included performed better than the submodels with variables from only one group included (models ABIOTIC and FOREST, Fig. 4). There was a difference between cross-validation results of winters with typical snow load conditions (2005-2017) and the 2017-2018 winter with exceptionally high snow loads. In the 2017-2018 winter the AUC values were also notably higher than in the results with full data or only years 2005-2017 and the ABIOTIC model with only abiotic predictors performed nearly as well as the full model (Fig. 4). In the cross-validation, the GLM and GAM models gave rather similar results. In general, GAM seemed to perform better for the ABIOTIC model and GLM for the FOREST model (Fig. 4).

**Figure 4.**
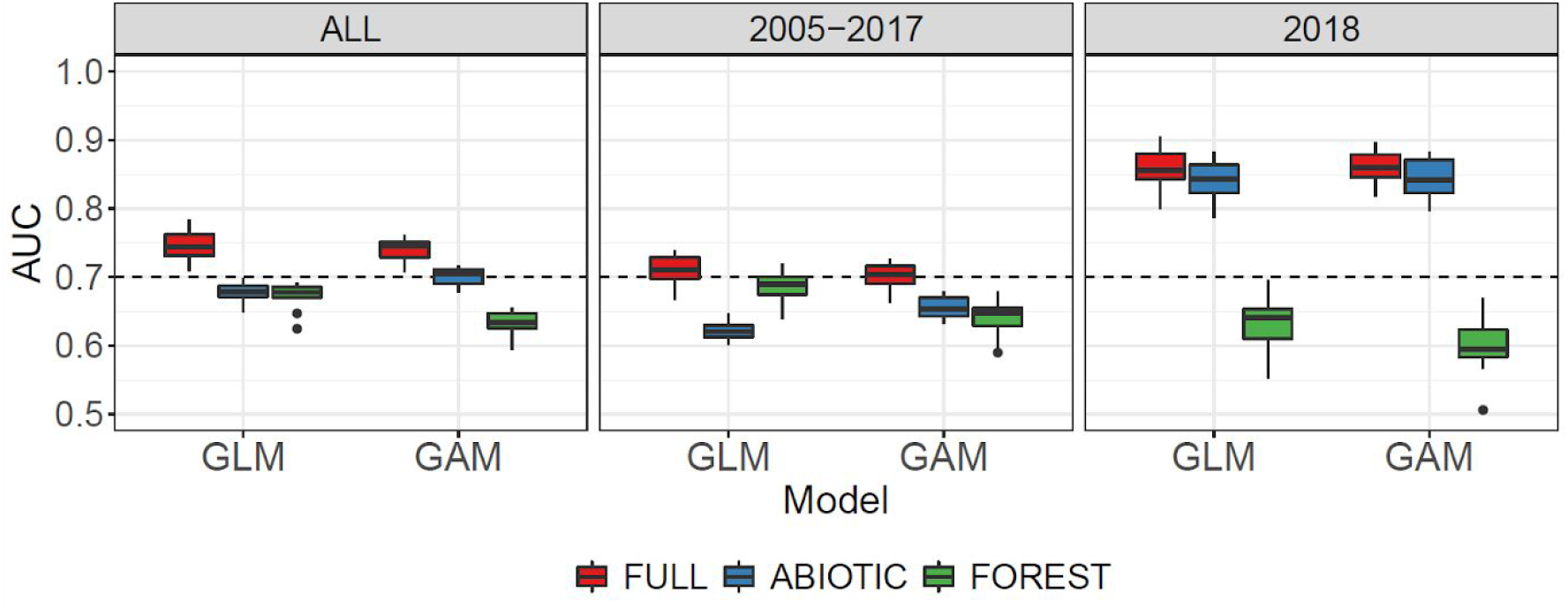
Cross-validation results for GLM and GAM with different predictor sets and for different time periods. Dash line shows the AUC=0.7 threshold for acceptable level of discrimination between cases and non-cases.

The snow damage probability maps predicted the highest snow damage risks in 2017-2018 near eastern border of the country (Fig. 5). The overall patterns in GLM and GAM maps were similar, with only minor differences. Testing the map with snow damage polygons showed that the model is able to predict damage probability on acceptable level also when GIS data is used for prediction instead of the field-measured NFI data (Fig. 6). Very high AUC values were obtained when the non-damage polygons were randomly sampled from the whole country (Fig. 6a) but also the test with non-damaged polygons sampled only from proximity of damaged polygons showed good ability of the model to identify the snow damaged polygons (Fig. 6b). The test showed quite similar results for the two modelling methods, though the map produced with the GAM model gained slightly better results (Fig. 6).

**Figure 5.**
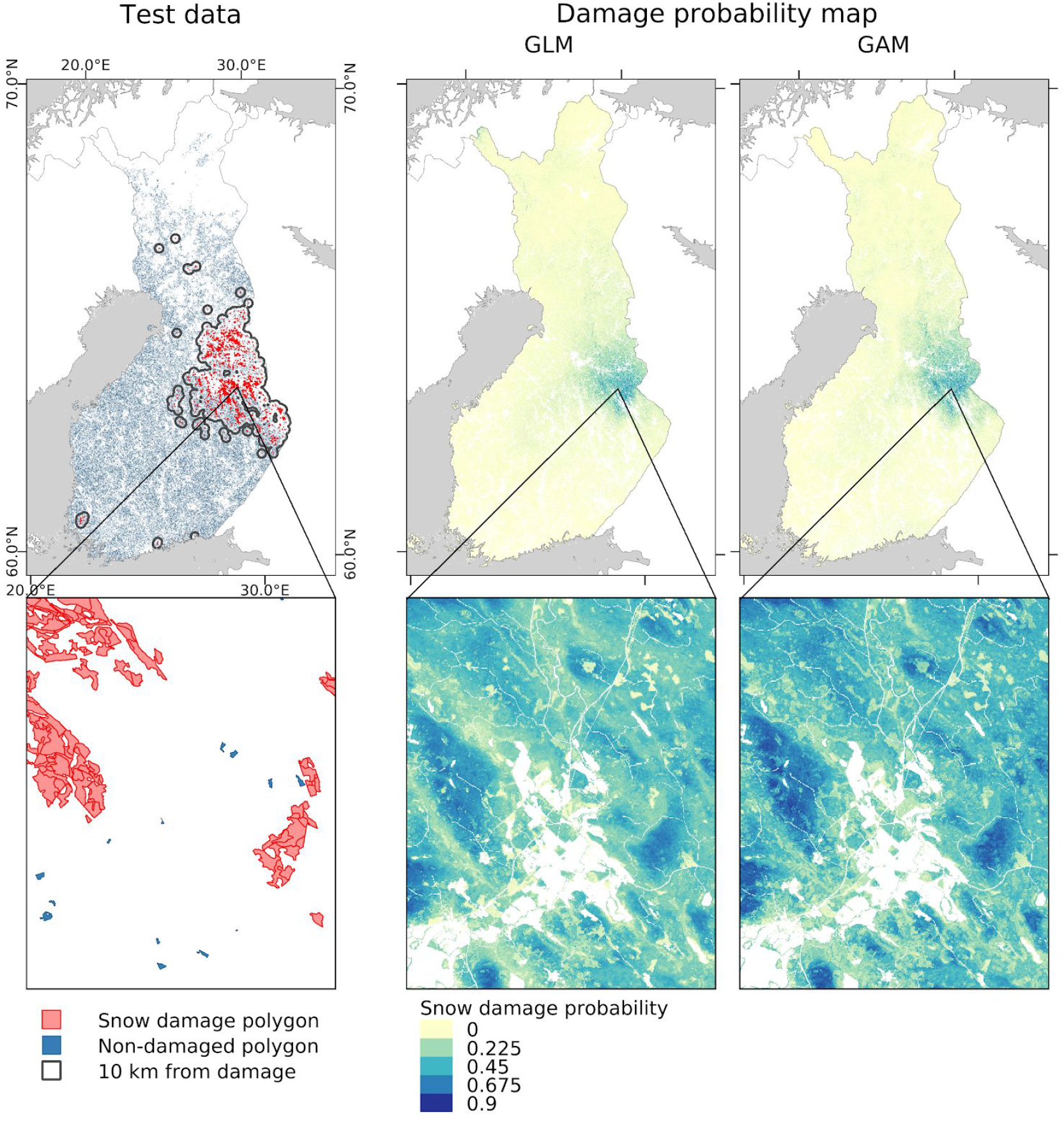
The Forest Centre data used for testing the snow damage probability maps, and the snow damage probability maps calculated with the snow load data from winter 2017-2018, using the full GLM and GAM models.

**Figure 6.**
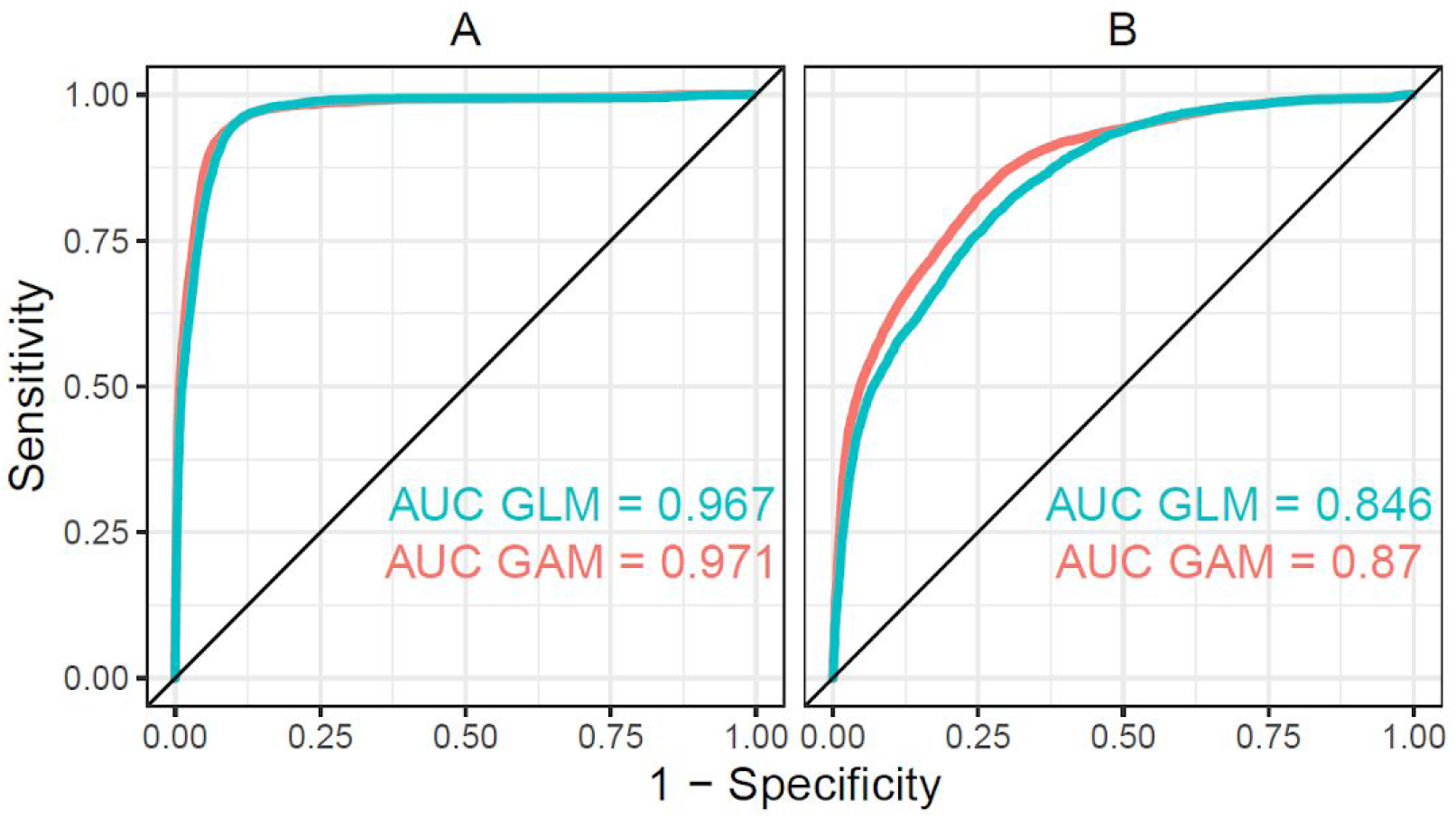
ROC curves and AUC values for the test of the snow damage probability map with the Forest Centre data: using non-damaged polygons from the whole country (A) and only considering non-damaged polygons within 10 km distance from snow damaged polygons (B).

## Discussion

We quantified the role of critical meteorological conditions to the snow damage risks by combining estimates of crown snow loads to the actual measurements of forest properties and snow damage from a large area in the boreal zone. The results showed that snow load becomes the dominating driver of damage during heavy snow years, but forest properties still improve the prediction of damage. During regular winters with typical snow packs, forest properties identify risk locations better than snow load and topographical information alone. Further, we demonstrated that the damage locations can be reliably pinpointed on heavy snow years at high resolution, which can be used to facilitate salvage logging and conservation planning. Moreover, the snow damage risk model can be applied with data of long-term snow load return-rates or projections of future snow loads, to generate risk estimates for the forest development scenarios under climate change.

The best predictions of snow damage probability were obtained when both abiotic variables (long term and recent snow load and topographic variables) and forest characteristics (species including an interaction with location in north boreal zone, dbh, basal area) were included in the model. By combining forest related predictors with snow load information from the winters preceding the NFI observations, our work extends further from many previous snow damage studies focusing solely on forest and site characteristics (Valinger and Fridman, 1999; Jalkanen and Mattila, 2000; Klopcic et al., 2009). While studies focusing on single snow damage events have been able to include both forest and snow information before (Hlásny et al., 2011), this is not the case for studies using long-term data from several damage events. In addition, the data describing snow load in the tree crown (Lehtonen et al., 2014), used in our analysis, provides a more realistic presentation of damage conditions compared to using information about snow depth, as used by, for example, Hlásny et al. (2011). On the other hand, our work offers potential improvements to meteorological estimations of snow load that assume a constant shape of tree crown in the calculation of snow load (Lehtonen et al., 2014, 2016), by incorporating detailed information about forest properties. This opens new possibilities for practical application possibilities, as increased accuracy in snow damage probability calculations can be attained when combining high-resolution forest data to the estimates formerly based only on simplified assumptions of the tree properties.

The exceptionally heavy snow load winter showed distinctively different patterns in our results compared to winters with lower snow load levels, as the model performed clearly better for the heavy snow load winter and the abiotic variables alone contributed for most of the model performance. This suggests that the processes of snow damage between heavy snow load winters and typical winter conditions have dissimilarities. It seems that during winters with low or moderate snow loads, snow disturbances only occur in the most vulnerable forests, emphasizing the importance of the forest predictors in these conditions. On the other hand, in winters with exceptionally high snow loads, damage can occur also on forests not as sensitive to snow damage, which is reflected in our results by the increased importance of the abiotic predictors. With lower snow loads forest properties drive the snow damage probability while their relative role diminishes when snow loads rise to exceptionally high levels. This interpretation is further supported by the effect of snow load in our results starting to strongly increase only after approx. 75 kg m^-2^, a level of crown snow load only rarely occurring in typical winter conditions (Fig. 1B). Uncertainties related to snow load data as well as the NFI damage observations may, at least partially, also play a role in the difference between typical and extreme snow load years. During a heavy snow load winter, the snow damage in forest is likely to be more clear and less likely by the NFI field team to be mistaken for wind damage. Meteorological estimation of snow loads may also be less uncertain when the snow loads are high.

The effects of abiotic and forest factors on snow damage probability in our final model were largely in line with the previous research. Increasing damage probability with elevation from sea level and with relative elevation from the surrounding terrain are backed by previous results (Makkonen and Ahti, 1995; Nykänen et al., 1997; Lehtonen et al., 2014). In accordance with literature a review by Nykänen (1997), damage probability in our results increased with basal area and decreased with stand average DBH. In our results, the effect of DBH was different for stands not dominated by pine or spruce (i.e., mainly birch dominated). Our results supported earlier research showing higher susceptibility to damage in coniferous versus deciduous trees and in Scots pine compared to Norway spruce (Suominen, 1963; Nykänen et al., 1997; Jalkanen and Mattila, 2000).

Our results reveal patterns suggesting adaptation of forests to high snow loads. First, in addition to the differences in damage probability between species, our results revealed geographical differences within Norway spruce, with spruce stands in the north boreal zone showing reduced probability of damage compared to spruce stands in the other parts of the country. The spruce trees in high latitude and altitude areas are known to have different crown morphology, with narrow crown shape reducing the accumulation of snow load on trees (Morgenstern 1996, Geburek et al. 2008). Our results show in practise how the morphological variation in the species leads to geographical variation in the predisposition of the trees to snow damage. Second, the negative effect of long-term snow load on the damage probability in the model suggests that forests in areas with historically higher snow loads are more resistant against snow damage. This effect was not species-specific but instead seems to affect stands regardless of the dominant species, as the interaction between long-term snow load and species was not statistically significant (S1). While the morphological differences may play a role also here, differences in forest structure in areas with high snow loads may also contribute in explaining this result, if the basal area and DBH included in the model are not sufficiently accounting for stand structure.

We did not find a statistically significant connection between thinnings and damage probability (results in S1). This finding contradicts previous research. According to literature review by Nykänen et al. (1997) trees in unthinned stands are more susceptible to snow damage and delayed thinning increases the snow damage risk. On the other hand, thinnings are found to temporarily increase the susceptibility of trees to snow damage, leading to higher damage risks during the first and second years after thinning (Suominen, 1963; Nykänen et al., 1997) and Wallentin and Nilsson (2014) found snow damage to be positively correlated with thinning invensity. We would not conclude from our results that forest management does not affect snow damage probability, instead the non-significance of the management effect is likely to be related to the insufficient detail of the used data. The main weakness in our analysis in regard to forest management is the imprecise definition of time between the damage and the management operation. As damage in our analysis had occurred within a time-window of five years before the NFI measurement, management operations could only be included if they had occurred more than five years ago. Otherwise it would not have been possible to differentiate between thinnings before the damage from ones occurring only after the damage. However, with this approach we are likely to lose the most sensitive period of one to two years after the thinning, when the damage sensitivity is the highest (Suominen, 1963; Nykänen et al., 1997). It is also worth noting that the forest variables included in the model (DBH and basal area) are strongly affected by management and, therefore, the effects of management are not completely excluded from the model.

Many previous studies have analyzed snow and wind damage together (Valinger and Fridman, 1999; Zhu et al., 2006; Suvanto et al., 2016; Díaz-Yáñez et al., 2019) as these two processes can act jointly in a damage event. For example, wind can more easily break trees with heavy snow load, or strong winds can either increase the snow accumulation or prevent the accumulation of snow on trees by shedding the snow from the branches (Solantie, 1994; Nykänen et al., 1997). However, while these processes can be related to each other, our results here and the previous results for wind damage (Suvanto et al., 2019) show that snow damage and wind damage affect different types of forest stands and have also spatially different occurrence patterns. While wind damage risk increases with tree height (Suvanto et al. 2019), snow damage is more typical in smaller trees, as shown in our results. In addition, snow damage can also occur with a minimal effect of wind (as in Hlásny et al. 2011) and wind disturbances often are not accompanied by snowfall.

The challenge in considering wind and snow separately in NFI data is in reliably identifying the cause of the damage in the field when field measurements are not targeting any specific damage event and stem breakages and uprooting can be related to either of the damage causes or their combined effects (Valinger and Fridman, 1999). For example, in southern Finland where heavy snow events are less common, snow damage may be mistakenly classified as wind damage, as those are more common in the region. The damage may have occurred already several years before the field measurement, making the correct identification of damage cause even harder. This adds uncertainty in the analysis and may also partly contribute to our results on why the model did not perform as well for the winters without heavy snow loads. Yet, while this uncertainty needs to be acknowledged we argue that, due to the differences in the two disturbance processes, it is beneficial to study damage caused by wind and by snow separately, whenever the used data makes this possible.

In our analysis, we did not differentiate between different snow damage types and, even though the used NFI data did contain some information on the damage type (see Table 1), stem breakage and uprooting were pooled in the same class, thus preventing us from analyzing them separately. This is a potential shortcoming of our analysis, as different damage dynamics may be behind stem breakage versus uprooting (Peltola et al., 1999; Zhu et al., 2006; Hlásny et al., 2011).

Logistic regression models (GLM) have long been the traditional method for modelling snow and other forest disturbances (Valinger and Fridman, 1999; Zhu et al., 2006; Suvanto et al., 2016; Díaz-Yáñez et al., 2019) whereas GAM provides more flexibility in modelling non-linear responses, as the relationship between continuous predictors and the response variable can be modelled with smoothing spline functions instead of the linear relationships used in the GLM. In our results, the comparison of the two statistical modelling methods showed rather similar results for the full model despite the method used. The GAM model performed better for the ABIOTIC model and the GLM for the FOREST model. This difference is likely to explain the better performance of the map based on the GAM model for the test data from winter 2017-2018, since this was a high snow load winter where the abiotic factors drove the damage probability. While the flexible spline functions in GAM increased the model performance in the case of the abiotic predictors, the traditionally used parametric models have some additional benefits in modelling forest disturbances, such as the ease of implementation of models in new applications and more straightforward interpretation of the models (Suvanto et al., 2019).

## Conclusions

In this study, we demonstrated the applicability of the damage probability mapping approach for snow disturbances, using NFI data in combination with GIS data layers, and tested the performance of the resulting map. The developed statistical model can be used to assess snow damage probability of forests, either in specific snow damage events by using observed snow load data or more generally by using data of long-term snow load return-rates or projections of future snow loads

Models with forest variables together with abiotic variables, including snow load, were found to perform better than models with predictors from only one of these variable groups. This was true especially for winters with typical snow load conditions, whereas the role of abiotic variables was emphasized in the heavy snow load winter. These results encourage combining snow load data with local and up-to date forest information, as increased accuracy in snow damage probability calculations can be attained when combining high-resolution forest data to the estimates formerly based only on simplified assumptions of the tree properties.

## Supporting information

Supplement S1: Model selection details.

## Acknowledgements

The research was funded from the project SÄÄTYÖ funded by the Ministry of Agriculture and Forestry of Finland. This project has received funding from the European Union’s Horizon 2020 research and innovation programme under the Marie Skłodowska-Curie grant agreement No 895158. We would like to thank the National Forest Inventory group in Luke for the NFI data we were able to use in the study. We acknowledge CSC – IT Center for Science, Finland, for computational resources.

## Supporting material

**S1.** Details of model selection

