## Supplement S1: Model selection details. for "Mapping the probability of forest snow disturbances in Finland"

#### 1. Summary

The model predictors were chosen based on:

1. existing understanding of snow damage dynamics,
2. availability of national extent GIS-data to be used for map prediction,
3. statistical significance of highest order terms in model, requiring significance on the level of $p < 0.001$.
4. improvement in AIC when comparing alternative models and
5. collinearity between predictors, generalized variation inflation factor (GVIF) < 4.

Table 1: Variables considered in model selection and reasons for exclusion from the final model (see numbering above).

| variable_name | description | included | reason |
|---|---|---|---|
| species_spruce | Dominant species spruce (ref.class=pine) | yes | – |
| species_other | Dominant species other (ref.class=pine) | yes | – |
| dbh | Diameter at breast height (DBH) | yes | – |
| basalarea | Basal area | yes | – |
| north | North boreal zone (0/1) | yes | – |
| snowload_longterm | Longterm average of max. snow load | yes | – |
| snowload | Snowload | yes | – |
| relelev_1km | Relative elevation | yes | – |
| dem | Altitude | yes | – |
| species_other:dbh | Interaction: Species, other x DBH | yes | – |
| species_spruce:north | Interaction: Species, spruce x North boreal zone | yes | – |
| species | Dominant species (ref.class=pine, 2=spruce, 3=other) | no | 3 |
| harvennus_yli5v | Thinning, > 5 yr | no | 3, 4 |
| ensiharvennus_yli5v | Precommercial thinning, >5 yr | no | 3, 4 |
| taimihoito_yli5v | Tending of seedling stands, >5 yr | no | 2, 3 |
| keskipit | Average tree height | no | 5, 3, 4 |
| kptyyp_2class | Site type | no | 3, 4 |
| species:basalarea | Interaction: Species x Basal area | no | 3 |
| species:dbh | Interaction: Species x DBH | no | 3 |
| snowload:north | Interaction: Snowload x North boreal zone | no | 3, 4 |
| north:dbh | Interaction: North x DBH | no | 3 |
| north:species | Interaction: North x Species | no | 3 |
| snowload_longterm:species | Interaction: Longterm snowload x Species | no | 3, 4 |
| n_species | Number of species | no | 2, 3 |
| max_sp_percent | Share of BA by dominant species | no | 3 |
| shannon_div | Shannon diversity index, calculated from species BA | no | 2, 3 |

For continuous variables with non-negative values log transformation was tested and kept in the model if produced a lower AIC.

Results for the variables not included in the final model are presented below.

#### 2. Thinning, > 5 years

Variable describes whether some type of thinning has been carried out at the stand more than five years ago (TRUE/FALSE).

Table 2: Model with variable "Thinning, > 5 years" (harvennus_yli5v)

| | Estimate | Std. Error | z value | Pr(>\|z\|) |
|---|---|---|---|---|
| (Intercept) | -7.6807848 | 0.1333161 | -57.6133418 | 0.0000000 |
| species2 | -0.3410196 | 0.0543925 | -6.2696075 | 0.0000000 |
| species3 | -0.4943257 | 0.0809380 | -6.1074608 | 0.0000000 |
| dbh | -0.0741110 | 0.0043993 | -16.8462490 | 0.0000000 |
| log(basalarea + 0.5) | 1.0774641 | 0.0478825 | 22.5022435 | 0.0000000 |
| harvennus_yli5vTRUE | 0.0087776 | 0.0848957 | 0.1033928 | 0.9176512 |
| snowload | 0.0290434 | 0.0008957 | 32.4264434 | 0.0000000 |
| relelev_1km | 0.0283242 | 0.0023644 | 11.9796878 | 0.0000000 |
| dem | 0.0038793 | 0.0003447 | 11.2549892 | 0.0000000 |

Table 3: AIC values for models with and without variable "Thinning, > 5 years" (harvennus_yli5v)

| | df | AIC |
|---|---|---|
| snowglm_with | 9 | 20760.39 |
| snowglm_without | 8 | 20758.40 |

#### 3. Precommercial thinning, > 5 years

Variable describes whether precommercial thinning has been carried out at the stand more than five years ago (TRUE/FALSE).

Table 4: Model with variable "Precommercial thinning, > 5 years" (ensiharvennus_yli5v)

| | Estimate | Std. Error | z value | Pr(>\|z\|) |
|---|---|---|---|---|
| (Intercept) | -7.6835003 | 0.1333067 | -57.6377688 | 0.0000000 |
| species2 | -0.3390768 | 0.0544250 | -6.2301692 | 0.0000000 |
| species3 | -0.4929056 | 0.0809425 | -6.0895756 | 0.0000000 |
| dbh | -0.0743611 | 0.0043538 | -17.0797438 | 0.0000000 |
| log(basalarea + 0.5) | 1.0781589 | 0.0478272 | 22.5427833 | 0.0000000 |
| ensiharvennus_yli5vTRUE | 0.0900984 | 0.0995052 | 0.9054644 | 0.3652194 |
| snowload | 0.0290292 | 0.0008957 | 32.4090946 | 0.0000000 |
| relelev_1km | 0.0282868 | 0.0023647 | 11.9622755 | 0.0000000 |

| | Estimate | Std. Error | z value | Pr(>\|z\|) |
|---|---|---|---|---|
| dem | 0.0038899 | 0.0003442 | 11.3026707 | 0.0000000 |

Table 5: AIC values for models with and without variable "Pre-commercial thinning, > 5 years" (ensiharvennus_yli5v)

| | df | AIC |
|---|---|---|
| snowglm_with_yli5v | 9 | 20759.6 |
| snowglm_without | 8 | 20758.4 |

### 4. Tending of seedling stands, > 5 years

Variable describes whether tending of seedling stand has been carried out at the stand more than five years ago (TRUE/FALSE).

Table 6: Model with variable "Tending of seedling stand, > 5 years" (taimihoito_yli5v)

| | Estimate | Std. Error | z value | Pr(>\|z\|) |
|---|---|---|---|---|
| (Intercept) | -7.6613039 | 0.1328692 | -57.660477 | 0.0000000 |
| species2 | -0.3375008 | 0.0543976 | -6.204327 | 0.0000000 |
| species3 | -0.4910299 | 0.0809519 | -6.065703 | 0.0000000 |
| dbh | -0.0766650 | 0.0044058 | -17.400812 | 0.0000000 |
| log(basalarea + 0.5) | 1.0913073 | 0.0478653 | 22.799540 | 0.0000000 |
| taimihoito_yli5vTRUE | -0.3319052 | 0.1062829 | -3.122846 | 0.0017911 |
| snowload | 0.0290211 | 0.0008961 | 32.386015 | 0.0000000 |
| relelev_1km | 0.0285544 | 0.0023641 | 12.078324 | 0.0000000 |
| dem | 0.0038746 | 0.0003434 | 11.281679 | 0.0000000 |

Table 7: AIC values for models with and without variable "Tending of seedling stand, > 5 years" (taimihoito_yli5v)

| | df | AIC |
|---|---|---|
| snowglm_with2 | 9 | 20749.76 |
| snowglm_without | 8 | 20758.40 |

### 5. Average tree height

The variable describes the average tree height of the stand (dm).

Table 8: Model with variable "Tree height" (keskipit)

| | Estimate | Std. Error | z value | Pr(>\|z\|) |
|---|---|---|---|---|
| (Intercept) | -7.6809409 | 0.1307901 | -58.727217 | 0 |
| species2 | -0.3790882 | 0.0547601 | -6.922711 | 0 |

| | Estimate | Std. Error | z value | Pr(>\|z\|) |
|---|---|---|---|---|
| species3 | -0.6434476 | 0.0835773 | -7.698831 | 0 |
| dbh | -0.1367818 | 0.0095640 | -14.301765 | 0 |
| log(basalarea + 0.5) | 0.9325585 | 0.0500466 | 18.633819 | 0 |
| keskipit | 0.0095572 | 0.0012630 | 7.567300 | 0 |
| snowload | 0.0290278 | 0.0008921 | 32.539084 | 0 |
| relelev_1km | 0.0280500 | 0.0023792 | 11.789502 | 0 |
| dem | 0.0048769 | 0.0003767 | 12.946961 | 0 |

Table 9: AIC values for models with and without variable "Tree height" (keskipit)

| | df | AIC |
|---|---|---|
| snowglm_with | 9 | 20702.31 |
| snowglm_without | 8 | 20758.40 |

Table 10: Variation inflation factors (GVIF) for model with variable "Tree height" (keskipit)

| | GVIF | Df | GVIF^(1/(2*Df)) |
|---|---|---|---|
| species | 1.119909 | 2 | 1.028716 |
| dbh | 8.290128 | 1 | 2.879258 |
| log(basalarea + 0.5) | 2.110775 | 1 | 1.452851 |
| keskipit | 9.902194 | 1 | 3.146775 |
| snowload | 1.180355 | 1 | 1.086442 |
| relelev_1km | 1.054602 | 1 | 1.026938 |
| dem | 1.476089 | 1 | 1.214944 |

Table 11: Model with variable "Tree height" (keskipit) while excluding DBH.

| | Estimate | Std. Error | z value | Pr(>\|z\|) |
|---|---|---|---|---|
| (Intercept) | -7.5311012 | 0.1318312 | -57.126832 | 0.00e+00 |
| species2 | -0.3465196 | 0.0545486 | -6.352496 | 0.00e+00 |
| species3 | -0.3309589 | 0.0803161 | -4.120706 | 3.78e-05 |
| log(basalarea + 0.5) | 0.9772503 | 0.0512775 | 19.058072 | 0.00e+00 |
| keskipit | -0.0069564 | 0.0005808 | -11.977487 | 0.00e+00 |
| snowload | 0.0292368 | 0.0008934 | 32.724019 | 0.00e+00 |
| relelev_1km | 0.0272636 | 0.0023391 | 11.655494 | 0.00e+00 |
| dem | 0.0027967 | 0.0003290 | 8.501453 | 0.00e+00 |

Table 12: Model with variable DBH while excluding "Tree height" (keskipit).

| | Estimate | Std. Error | z value | Pr(>\|z\|) |
|---|---|---|---|---|
| (Intercept) | -7.6801166 | 0.1331415 | -57.683872 | 0 |

| | Estimate | Std. Error | z value | Pr(>\|z\|) |
|---|---|---|---|---|
| species2 | -0.3411606 | 0.0543756 | -6.274153 | 0 |
| species3 | -0.4944811 | 0.0809239 | -6.110442 | 0 |
| dbh | -0.0740317 | 0.0043307 | -17.094747 | 0 |
| log(basalarea + 0.5) | 1.0771217 | 0.0477597 | 22.552942 | 0 |
| snowload | 0.0290449 | 0.0008956 | 32.431754 | 0 |
| relelev_1km | 0.0283292 | 0.0023638 | 11.984654 | 0 |
| dem | 0.0038766 | 0.0003437 | 11.280454 | 0 |

Table 13: AIC values for models with either Tree height (keskipit) or DBH included.

| | df | AIC |
|---|---|---|
| snowglm_keskipit | 8 | 20928.56 |
| snowglm_dbh | 8 | 20758.40 |

Table 14: Variation inflation factors (GVIF) for model with variable Tree height" (keskipit) while excluding DBH.

| | GVIF | Df | GVIF^(1/(2*Df)) |
|---|---|---|---|
| species | 1.035841 | 2 | 1.008842 |
| log(basalarea + 0.5) | 2.018151 | 1 | 1.420616 |
| keskipit | 2.034260 | 1 | 1.426275 |
| snowload | 1.186678 | 1 | 1.089348 |
| relelev_1km | 1.062092 | 1 | 1.030578 |
| dem | 1.281239 | 1 | 1.131918 |

Table 15: Variation inflation factors (GVIF) for model with variable DBH while excluding “Tree height” (keskipit).

| | GVIF | Df | GVIF^(1/(2*Df)) |
|---|---|---|---|
| species | 1.042045 | 2 | 1.010350 |
| dbh | 1.736970 | 1 | 1.317941 |
| log(basalarea + 0.5) | 1.774385 | 1 | 1.332061 |
| snowload | 1.182050 | 1 | 1.087221 |
| relelev_1km | 1.058808 | 1 | 1.028984 |
| dem | 1.295870 | 1 | 1.138363 |

### 6. Site type

Site type “FERTILE” or “POOR”, reclassification of site type classes in the Finnish NFI (classes 1-3 = FERTILE, classes > 3 = POOR).

Table 16: Model with variable "Site type" (kptyyp_2class)

| | Estimate | Std. Error | z value | Pr(>\|z\|) |
|---|---|---|---|---|
| (Intercept) | -7.6708986 | 0.1411311 | -54.3530000 | 0.0000000 |
| species2 | -0.3453836 | 0.0584837 | -5.9056382 | 0.0000000 |
| species3 | -0.4986814 | 0.0837127 | -5.9570607 | 0.0000000 |
| dbh | -0.0740416 | 0.0043304 | -17.0980150 | 0.0000000 |
| log(basalarea + 0.5) | 1.0757593 | 0.0482355 | 22.3022110 | 0.0000000 |
| kptyyp_2classPOOR | -0.0096050 | 0.0490838 | -0.1956861 | 0.8448558 |
| snowload | 0.0290470 | 0.0008957 | 32.4299531 | 0.0000000 |
| relelev_1km | 0.0282998 | 0.0023685 | 11.9482003 | 0.0000000 |
| dem | 0.0038734 | 0.0003440 | 11.2604828 | 0.0000000 |

Table 17: AIC values for models with and without variable "Site type" (kptyyp_2class)

| | df | AIC |
|---|---|---|
| snowglm_with | 9 | 20760.36 |
| snowglm_without | 8 | 20758.40 |

### 7. Interactions: Species x DBH or Basal area

Interactions between two predictors:

- Species x DBH
- Species x Basal area

Table 18: Model with interaction Species x DBH

| | Estimate | Std. Error | z value | Pr(>\|z\|) |
|---|---|---|---|---|
| (Intercept) | -7.6890780 | 0.1394996 | -55.119006 | 0.0000000 |
| species2 | -0.5927490 | 0.1407292 | -4.211982 | 0.0000253 |
| species3 | 0.6395325 | 0.2183468 | 2.928975 | 0.0034008 |
| dbh | -0.0746229 | 0.0050388 | -14.809710 | 0.0000000 |
| log(basalarea + 0.5) | 1.0833878 | 0.0479459 | 22.596030 | 0.0000000 |
| snowload | 0.0291418 | 0.0008985 | 32.432293 | 0.0000000 |
| relelev_1km | 0.0281579 | 0.0023666 | 11.897829 | 0.0000000 |
| dem | 0.0038479 | 0.0003437 | 11.195885 | 0.0000000 |
| species2:dbh | 0.0139089 | 0.0073270 | 1.898313 | 0.0576548 |
| species3:dbh | -0.0877992 | 0.0165816 | -5.294979 | 0.0000001 |

Table 19: Model with interaction Species x Basal area

| | Estimate | Std. Error | z value | Pr(>\|z\|) |
|---|---|---|---|---|
| (Intercept) | -7.9718471 | 0.1597659 | -49.8970604 | 0.0000000 |
| species2 | -0.2182899 | 0.2551237 | -0.8556238 | 0.3922059 |
| species3 | 1.8375482 | 0.2872057 | 6.3980212 | 0.0000000 |

| | Estimate | Std. Error | z value | Pr(>\|z\|) |
|---|---|---|---|---|
| dbh | -0.0761704 | 0.0043713 | -17.4251090 | 0.0000000 |
| log(basalarea + 0.5) | 1.1856997 | 0.0562248 | 21.0885598 | 0.0000000 |
| snowload | 0.0291187 | 0.0008965 | 32.4804245 | 0.0000000 |
| relelev_1km | 0.0281458 | 0.0023636 | 11.9079099 | 0.0000000 |
| dem | 0.0039801 | 0.0003446 | 11.5489101 | 0.0000000 |
| species2:log(basalarea + 0.5) | -0.0450562 | 0.0856941 | -0.5257793 | 0.5990416 |
| species3:log(basalarea + 0.5) | -0.8526833 | 0.1055349 | -8.0796311 | 0.0000000 |

Table 20: AIC values for models with interaction Species x BA, with interaction Species x DBH and without interactions.

| | df | AIC |
|---|---|---|
| snowglm_withBA | 10 | 20710.78 |
| snowglm_withDBH | 10 | 20725.53 |
| snowglm_without | 8 | 20758.40 |

### 8. North boreal zone and interactions

Adding the north boreal zone ('north') into the model.

#### 8.1 North

Table 21: Model with variable 'north' describing if plot located in north boreal zone.

| | Estimate | Std. Error | z value | Pr(>\|z\|) |
|---|---|---|---|---|
| (Intercept) | -7.7207920 | 0.1329452 | -58.075010 | 0.0000000 |
| species2 | -0.3473889 | 0.0544970 | -6.374455 | 0.0000000 |
| species3 | -0.4952526 | 0.0809362 | -6.119046 | 0.0000000 |
| dbh | -0.0737315 | 0.0043357 | -17.005868 | 0.0000000 |
| log(basalarea + 0.5) | 1.0816260 | 0.0482313 | 22.425796 | 0.0000000 |
| snowload | 0.0289589 | 0.0008986 | 32.227654 | 0.0000000 |
| relelev_1km | 0.0280042 | 0.0023811 | 11.761227 | 0.0000000 |
| dem | 0.0041480 | 0.0004152 | 9.990217 | 0.0000000 |
| north | -0.0739752 | 0.0663517 | -1.114895 | 0.2648954 |

Table 22: AIC values for models with and without variable 'north'.

| | df | AIC |
|---|---|---|
| snowglm_without | 8 | 20757.61 |
| snowglm_with | 9 | 20758.36 |

##### 8.2 North x snowload

Testing interaction with snowload, as northern trees, adapted to heavy snowloads, could response to snowload differently.

Table 23: Model with interaction north x snowload

| | Estimate | Std. Error | z value | Pr(>\|z\|) |
|---|---|---|---|---|
| (Intercept) | -7.7048499 | 0.1335721 | -57.683082 | 0.0000000 |
| species2 | -0.3489004 | 0.0544964 | -6.402259 | 0.0000000 |
| species3 | -0.4957242 | 0.0809140 | -6.126555 | 0.0000000 |
| dbh | -0.0738949 | 0.0043380 | -17.034431 | 0.0000000 |
| log(basalarea + 0.5) | 1.0818972 | 0.0482135 | 22.439695 | 0.0000000 |
| snowload | 0.0286110 | 0.0009489 | 30.152055 | 0.0000000 |
| relelev_1km | 0.0279844 | 0.0023832 | 11.742412 | 0.0000000 |
| dem | 0.0041992 | 0.0004177 | 10.052605 | 0.0000000 |
| north | -0.3232467 | 0.2235972 | -1.445665 | 0.1482712 |
| snowload:north | 0.0034789 | 0.0029661 | 1.172865 | 0.2408501 |

Table 24: AIC values for models with and without interaction north x species

| | df | AIC |
|---|---|---|
| snowglm_without | 8 | 20757.61 |
| snowglm_with | 10 | 20758.99 |

##### 8.3 North x DBH

Testing interaction with DBH, as northern trees, adapted to heavy snowloads, could have different pattern of DBH effect on damage probability.

Table 25: Model with interaction north x DBH

| | Estimate | Std. Error | z value | Pr(>\|z\|) |
|---|---|---|---|---|
| (Intercept) | -7.6770374 | 0.1354296 | -56.686537 | 0.0000000 |
| species2 | -0.3506124 | 0.0544991 | -6.433363 | 0.0000000 |
| species3 | -0.4980986 | 0.0809638 | -6.152114 | 0.0000000 |
| dbh | -0.0763828 | 0.0046608 | -16.388475 | 0.0000000 |
| log(basalarea + 0.5) | 1.0844146 | 0.0482347 | 22.482029 | 0.0000000 |
| snowload | 0.0289710 | 0.0008988 | 32.234397 | 0.0000000 |
| relelev_1km | 0.0279410 | 0.0023786 | 11.746599 | 0.0000000 |
| dem | 0.0040873 | 0.0004170 | 9.801631 | 0.0000000 |
| north | -0.2778253 | 0.1465753 | -1.895444 | 0.0580336 |
| dbh:north | 0.0131626 | 0.0083709 | 1.572416 | 0.1158540 |

Table 26: AIC values for models with and without interaction north x DBH

| | df | AIC |
|---|---|---|
| snowglm_without | 8 | 20757.61 |
| snowglm_with | 10 | 20757.88 |

##### 8.4 North x Species

Testing interaction with DBH, as different adaptations to northern conditions are known between species, morphological adaptation being especially clear in Norway spruce.

Table 27: Model with interaction north x species

| | Estimate | Std. Error | z value | Pr(>\|z\|) |
|---|---|---|---|---|
| (Intercept) | -7.7690642 | 0.1331775 | -58.3361554 | 0.0000000 |
| species2 | -0.2531935 | 0.0578732 | -4.3749697 | 0.0000121 |
| species3 | -0.4178832 | 0.0851977 | -4.9048672 | 0.0000009 |
| dbh | -0.0727090 | 0.0043321 | -16.7838777 | 0.0000000 |
| log(basalarea + 0.5) | 1.0756412 | 0.0481114 | 22.3572976 | 0.0000000 |
| snowload | 0.0289123 | 0.0008991 | 32.1558917 | 0.0000000 |
| relelev_1km | 0.0273759 | 0.0023876 | 11.4657034 | 0.0000000 |
| dem | 0.0043324 | 0.0004181 | 10.3625481 | 0.0000000 |
| north | 0.0213918 | 0.0687567 | 0.3111231 | 0.7557071 |
| species2:north | -0.7517200 | 0.1862165 | -4.0368058 | 0.0000542 |
| species3:north | -0.5686186 | 0.2803981 | -2.0278977 | 0.0425707 |

Table 28: AIC values for models with and without interaction north x DBH

| | df | AIC |
|---|---|---|
| snowglm_without | 8 | 20757.61 |
| snowglm_with | 11 | 20740.17 |

#### 9. Long term snowload x species

Testing the interaction between long term snowload (15 years average of winter max) and species, as Norway spruce shows clearer morphological differentiation with latitude and could therefore show species-specific reponse to long-term snowload.

Table 29: Model with interaction Long term snowload x species

| | Estimate | Std. Error | z value | Pr(>\|z\|) |
|---|---|---|---|---|
| (Intercept) | -7.2849719 | 0.1841717 | -39.5553273 | 0.0000000 |
| species_spruce | 0.1068927 | 0.2860188 | 0.3737262 | 0.7086080 |
| species_other | 0.7234707 | 0.4648012 | 1.5565164 | 0.1195854 |
| dbh | -0.0732843 | 0.0043957 | -16.6719259 | 0.0000000 |
| log(basalarea + 0.5) | 1.1190323 | 0.0484103 | 23.1155616 | 0.0000000 |

| | Estimate | Std. Error | z value | Pr(>\|z\|) |
|---|---|---|---|---|
| snowload_longterm | -0.0243462 | 0.0048049 | -5.0669548 | 0.0000004 |
| snowload | 0.0325340 | 0.0010532 | 30.8897068 | 0.0000000 |
| relelev_1km | 0.0270099 | 0.0023912 | 11.2953546 | 0.0000000 |
| dem | 0.0050485 | 0.0003924 | 12.8652716 | 0.0000000 |
| species_other:dbh | -0.0920792 | 0.0163444 | -5.6336874 | 0.0000000 |
| species_spruce:snowload_longterm | -0.0130581 | 0.0076964 | -1.6966466 | 0.0897635 |
| species_other:snowload_longterm | -0.0005436 | 0.0111923 | -0.0485689 | 0.9612629 |

Table 30: AIC values for models with and without interaction Long term snowload x species

| | df | AIC |
|---|---|---|
| snowglm_without | 10 | 20692.53 |
| snowglm_with | 12 | 20693.62 |

### 10. Species composition

The effect of species composition was studied with three different variables:

- number of species
- BA proportion of most abundant species (most abundant here meaning the species with largest basal area)
- Shannon diversity index (calculated from % of basal area)

Table 31: Model with number of tree species (n_species).

| | Estimate | Std. Error | z value | Pr(>\|z\|) |
|---|---|---|---|---|
| (Intercept) | -7.0574239 | 0.1313806 | -53.7173884 | 0.0000000 |
| species_spruce | -0.2681443 | 0.0586793 | -4.5696554 | 0.0000049 |
| species_other | 0.7433953 | 0.2149934 | 3.4577594 | 0.0005447 |
| dbh | -0.0728983 | 0.0043807 | -16.6406130 | 0.0000000 |
| log(basalarea + 0.5) | 1.1190169 | 0.0489719 | 22.8501796 | 0.0000000 |
| log(n_species) | -0.1060911 | 0.0467952 | -2.2671358 | 0.0233819 |
| snowload_anomaly | 0.0330034 | 0.0009908 | 33.3092338 | 0.0000000 |
| relelev_1km | 0.0256751 | 0.0024017 | 10.6904942 | 0.0000000 |
| dem | 0.0059132 | 0.0004028 | 14.6797453 | 0.0000000 |
| north | -0.0522483 | 0.0690748 | -0.7564027 | 0.4494078 |
| species_other:dbh | -0.0922254 | 0.0163765 | -5.6315877 | 0.0000000 |
| species_spruce:north | -0.7439038 | 0.1860639 | -3.9981099 | 0.0000639 |

Table 32: Model with proportion of BA covered by the most abundant species (max_sp_percent).

| | Estimate | Std. Error | z value | Pr(>\|z\|) |
|---|---|---|---|---|
| (Intercept) | -7.3518540 | 0.1749239 | -42.0288627 | 0.0000000 |

| | Estimate | Std. Error | z value | Pr(>\|z\|) |
|---|---|---|---|---|
| species_spruce | -0.2660797 | 0.0587368 | -4.5300340 | 0.0000059 |
| species_other | 0.7380216 | 0.2143258 | 3.4434572 | 0.0005743 |
| dbh | -0.0728918 | 0.0043809 | -16.6385639 | 0.0000000 |
| log(basalarea + 0.5) | 1.1140774 | 0.0488026 | 22.8282622 | 0.0000000 |
| max_sp_percent | 0.2925138 | 0.1239580 | 2.3597807 | 0.0182857 |
| snowload_anomaly | 0.0329717 | 0.0009897 | 33.3145282 | 0.0000000 |
| relelev_1km | 0.0255505 | 0.0024013 | 10.6401676 | 0.0000000 |
| dem | 0.0059055 | 0.0004028 | 14.6607314 | 0.0000000 |
| north | -0.0477269 | 0.0689040 | -0.6926569 | 0.4885249 |
| species_other:dbh | -0.0909531 | 0.0163410 | -5.5659568 | 0.0000000 |
| species_spruce:north | -0.7336440 | 0.1862241 | -3.9395766 | 0.0000816 |

Table 33: Model with Shannon diversity index (shannon_div).

| | Estimate | Std. Error | z value | Pr(>\|z\|) |
|---|---|---|---|---|
| (Intercept) | -7.1817792 | 0.1360826 | -52.7751643 | 0.0000000 |
| species_spruce | -0.2643695 | 0.0585866 | -4.5124552 | 0.0000064 |
| species_other | 0.7476963 | 0.2146353 | 3.4835655 | 0.0004948 |
| dbh | -0.0727750 | 0.0043761 | -16.6300508 | 0.0000000 |
| log(basalarea + 0.5) | 1.1190954 | 0.0488620 | 22.9031634 | 0.0000000 |
| log(shannon_div + 1e-04) | -0.0157360 | 0.0054827 | -2.8701187 | 0.0041032 |
| snowload_anomaly | 0.0330205 | 0.0009905 | 33.3364265 | 0.0000000 |
| relelev_1km | 0.0255899 | 0.0024028 | 10.6502298 | 0.0000000 |
| dem | 0.0059209 | 0.0004030 | 14.6917770 | 0.0000000 |
| north | -0.0536816 | 0.0690004 | -0.7779896 | 0.4365751 |
| species_other:dbh | -0.0922571 | 0.0163489 | -5.6430204 | 0.0000000 |
| species_spruce:north | -0.7316969 | 0.1861833 | -3.9299818 | 0.0000850 |

Table 34: AIC values of models with species community variables.

| | df | AIC |
|---|---|---|
| snowglm_without | 11 | 20674.88 |
| snowglm_n_species | 12 | 20671.75 |
| snowglm_max_sp_percent | 12 | 20671.25 |
| snowglm_shannon | 12 | 20668.74 |
